# Quality of life and depressive symptoms among Peruvian university students during the COVID-19 pandemic

**DOI:** 10.1101/2020.12.04.411330

**Authors:** Joel Figueroa-Quiñones, Miguel Ipanaqué-Zapata, Daniel Machay-Pak, Juan Rodríguez-Ruiz

## Abstract

**Objectives:** Characterize the quality of life and depressive symptoms of university students in Peru during the COVID-19 pandemic and to determine the associated factors.

**Methods:** Multi-centre study in 1634 university students recruited by convenience sampling. Quality of life (QoL) was assessed with the European Quality of Life-5 Dimensions at three levels (EQ-5D-3L) and depressive symptoms with the Patient Health Questionarie-9 (PHQ-9). To evaluate factors associated with QoL and depressive symptoms, linear and adjusted regressions were used, with robust variance reporting coefficients (β).

**Results:** The percentage of participants most affected by QoL dimension was: anxiety/depression (47.2%) and pain/discomfort (35.6%). Regarding the Visual Analog Scale (EQ-VAS) of QoL, the score was 76.0 + 25.6. Those who had family economic decline during quarantine (β=-3.4, IC95%=-6.5 to −0.3) or family with chronic diseases (β=-3.7, IC95%=-6.1 to −1.4) presented significantly lower scores in their QoL. Regarding depressive symptomatology, the university students reported a moderate to severe level (28.9%). A higher risk of depressive symptoms was found in residents of Ayacucho (β=0.8, IC95%=0.1 to 1.5), those who were released from quarantine (β=0.7, IC95%=0.2 to 1.2) and those who had a family member with chronic disease (β=1.5, IC95%=1.0 to 2.1).

**Conclusions:** Almost half and one third of participants reported anxiety/depression, and pain/discomfort in their QoL respectively. Nearly a third presented moderate and severe depressive symptoms. The deterioration of QoL was worse in those who had a decrease in income and a family member with chronic illness. The presence of depressive symptoms was found in students in Ayacucho, those who left home during quarantine and those who had a family member with chronic diseases.

## Introduction

Coronavirus disease (COVID-19) classified as a pandemic by the World Health Organization (WHO) is a public health problem that has affected the entire world population. COVID-19 presents signs and symptoms similar to those of a common cold, but its complication in patients at risk can be fatal (1). Worldwide, COVID-19 has infected more than 50 million people and caused more than 1 million deaths (2). In Latin America over 12 million cases of COVID-19 and 400,000 deaths were recorded up to November 2020, making it one of the regions most affected (3).

The measures to control the spread of COVID-19, such as confinement, have brought about changes in the routine of life of the population that could affect their mental health (4, 5). A recent review study reported prevalences of anxious and depressive symptoms in the population in ranges of 6-50% and 14-48% respectively (6). Some factors have been associated with a decline in mental health during the COVID-19 pandemic. For example, people with low economic status, women, and the unemployed have experienced greater mental health problems (7,8). In contrast, a study in China found that the population with depressive symptoms reported a lower quality of life (9). To date, few studies have reported on the COVID-19 pandemic and depressive symptoms and quality of life (QoL) in students despite the fact that the pandemic has severely affected various parts of the world

The rapid transmission of COVID-19 led to the suspension of face-to-face academic activities, affecting 91.3% of the world’s student population, who had to adapt quickly to non-face-to-face education (10). However, this abrupt transition has led to increased stress, insomnia, anxiety and depression in students, so that their quality of life and mental health may have deteriorated (11–13). Peru reported the first case of COVID-19 on 06 March 2020 and 10 days later a state of national emergency and compulsory social isolation (confinement) was declared (14), but the virus spread rapidly throughout the country. A total of 950, 557 people were infected by COVID-19 and 35,641 deaths were recorded in the country as of 24 November 2020 (15).

The confinement affected the economy of the families and brought anguish to the students of Peru, due to the economic difficulties and digital gaps (connectivity, computers), which caused the government to implement economic policies to the Peruvian families and the education sector such as economic bonds, educational credits, connectivity and scholarships (16). However, there are currently few studies on the situation of QoL and depressive symptoms in university students in the current pandemic situation despite their vulnerable condition (17, 18), which makes it difficult to implement programmes and interventions to improve mental health and quality of life in the face of the COVID-19 public health emergency.

Therefore, this study sought to characterize the depressive symptoms and quality of life of university students during the COVID-19 pandemic as well as to determine the association of depression with quality of life in university students in Peru during the COVID-19 pandemic.

## Materials and methods

### Research design and context

Multi-centre cross-sectional analytical study in four departments of Peru (Ancash, Lima, Ayacucho and Piura) of a private university in Peru in the period July-August 2020.

Peru is located in the central part of South America near the Pacific Ocean. Furthermore, it has an extension of 1’285 215.6 km2 and is divided into 25 departments with an estimated population of 32’636 000 habitants for the present year.

### Participants

The participants in the study were 1634 university students from four departments of Peru (Ancash, Piura, Lima and Ayacucho), which were obtained from a non-probabilistic sampling for convenience. Students duly enrolled in the private university of Peru, those who accepted to participate voluntarily in the study or those who managed to complete the survey were included.

### Procedures

The information collected for the study was through the application of an online survey using SurveyMonkey’s virtual platform (19). This survey was reviewed by all the authors and the SurveyMonkey administrator before it was published. The publication of the survey was hosted and promoted by emails and study groups of university students and they were informed of the objective and nature of the study, ensuring their anonymity and confidentiality of the data. Finally, the students who decided to participate in the study and entered the online survey were informed in detail and asked to accept the informed consent, to later conduct the survey. An email and WhatsApp were opened by the authors of the study to answer any questions from participants.

The study information was collected through a spreadsheet, maintaining anonymity with no way of linking the questionnaires to the participants, and was protected through computer storage and password access to the server only.

### Variables

The main variables of the study were quality of life (QoL) and depressive symptoms in university students in four sites in Peru. The QoL variable was measured through the European Quality of Life-5 Dimensions questionnaire in three levels (EQ-5D-3L) (20), comprised of 5 items that respond to 5 dimensions (Mobility, Personal Care, Activities of Daily Living, Pain and Depression or Anxiety), each dimension has 3 levels of response of ordinal type according to the presence of each dimension: absence, moderate presence and severe presence. Likewise, it presents an analogous visual scale (EQ-VAS) that consists of reporting the current state of life within a range of 0 (worst state) to 100 (best state), being this last one the quality of life indicator for the present study. The instrument is used worldwide in both English and Spanish; there are also previous studies carried out in Peru (21, 22).

With respect to the depressive symptoms variable, the Patient Health Questionarie-9 (PHQ-9) was used. It is a self-administered questionnaire of 9 items with 4 levels of response according to the frequency of depressive symptoms within the last 2 weeks: nothing, several days, more than half the days and almost every day. This questionnaire is validated in the Peruvian population presenting adequate indicators of psychometric properties and reliability (α = 0.87) (23). The summed score is within the range 0 to 27 points. Likewise, depressive symptoms are reported according to levels of severity: minimal (0-4), mild (5-9), moderate (10-14), moderately severe (15–19) and severe (20–27) (24–26).

The covariates that participated in the study were: age (in tertiles), sex (male or female), department of residence (Ancash, Ayacucho, Lima and Piura), marital status (single/separate/widowed/divorced and married/cohabitant), occupation (studies and works, only work), left home during quarantine (no and yes), decreased family income in quarantine (no and yes), lives alone (no and yes) and family with chronic illness (no and yes).

### Data analysis

Descriptive analysis was presented through measures of central tendency and dispersion (for numerical variables) and absolute frequencies (for categorical variables). Raw and adjusted robust variance linear regression models reporting coefficients (β) with their 95% confidence intervals (CI95%) were performed to assess the factors associated with the EQ-VAS (quality of life) and PHQ-9 (depressive symptoms) scores. In all cases, the variables that obtained a p<0.20 in the raw model were included in the adjusted model. The analyses were performed in the statistical software Stata v15.0 (27).

### Ethical principles

This study was approved by the Institutional Committee on Research Ethics (CIEI) at Los Angeles Catholic University in Chimbote. Participants voluntarily accepted an informed consent within the survey to be part of the study. The principles of justice, confidentiality and autonomy that are conducive to human studies were also applied and respected (28).

## Results

### Characteristics of respondents

Initially, 1825 participants were recruited, of which 65 did not agree to participate in the study and 126 failed to complete the entire survey; therefore, only 1634 (89.5%) university students participated in the study. The participants included were distributed by department of residence: 712 (43.6%) were from Ayacucho, 347 (21.2%) from Ancash, 342 (20.9%) from Piura, and 233 (14.3%) from Lima (Table 1).

**Table 1.**
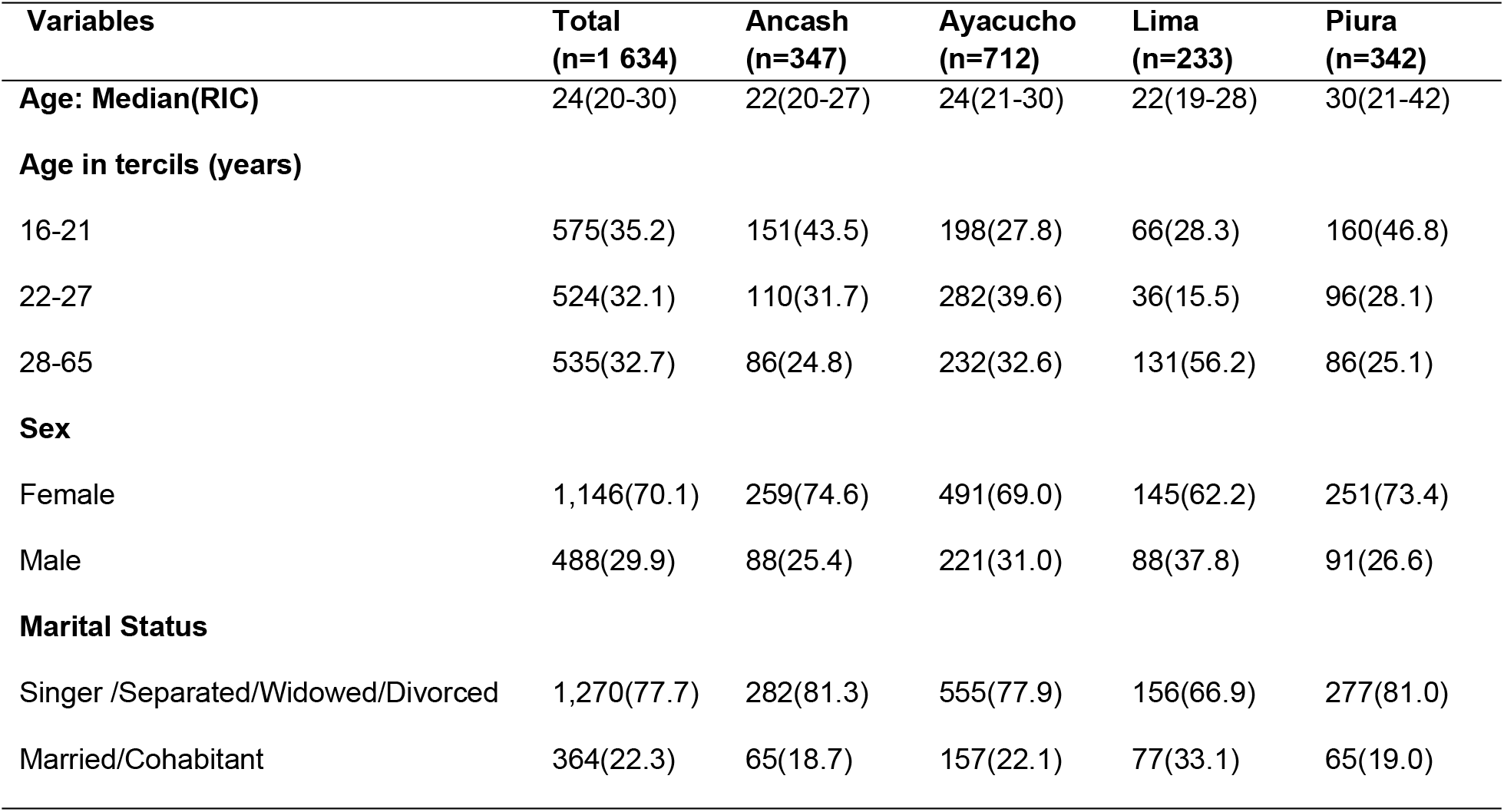
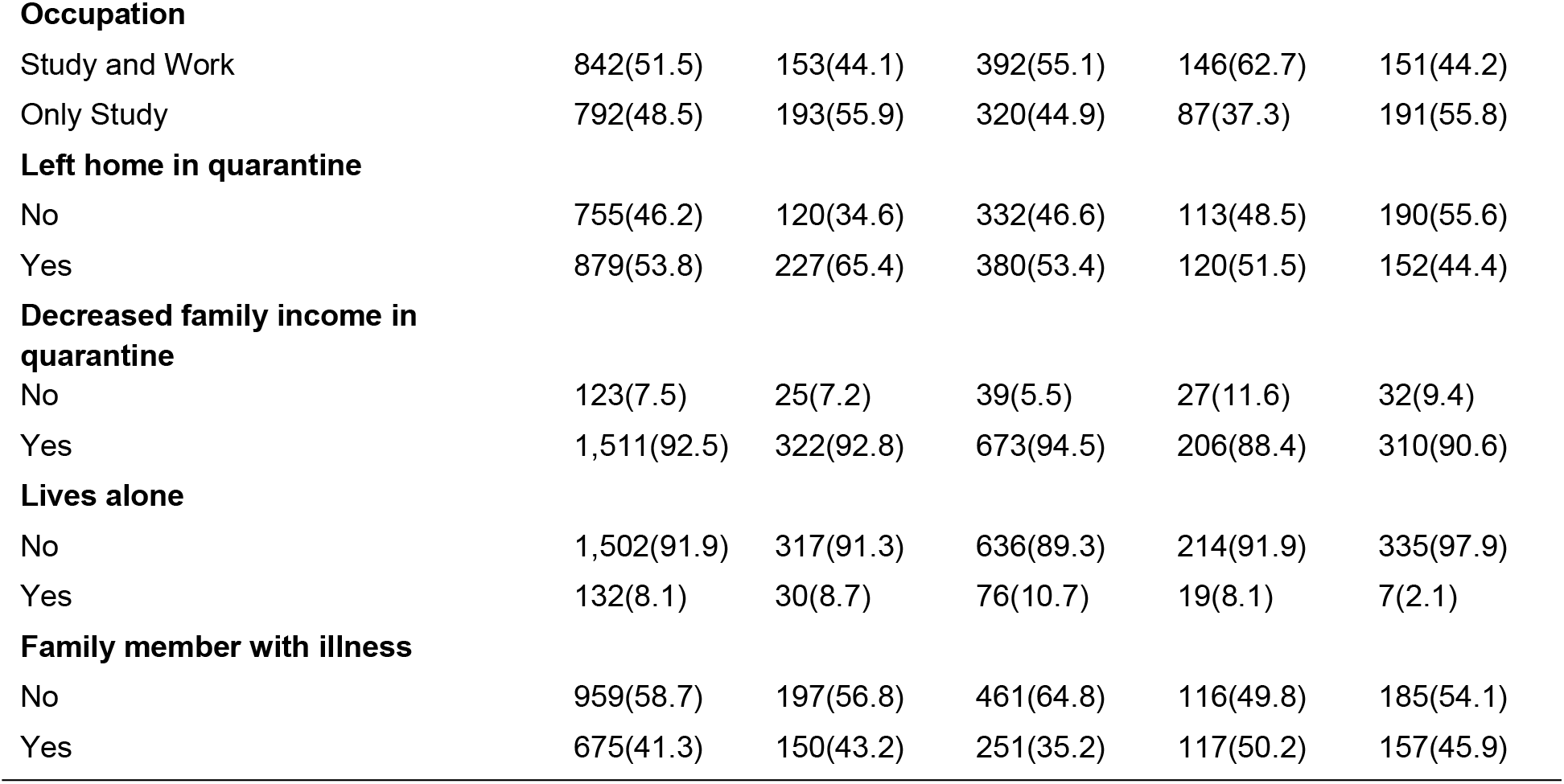
Characteristics of the populations assessed

The participants had a median age of 24 years (interquartile range: 20 to 30 years), 1146 (70.1%) were women, 1270 (77.7%) reported being within the marital status single, separated, widowed or divorced. University students studying and working at the same time were 842 (51.5%) with a significant prevalence in Lima. 879 (53.8%) of university students reported leaving home during quarantine, 1511 (92.5%) had a decrease in family income, 1502 (91.9%) reported not living alone and 959 (58.69%) had a family member with a chronic illness (Table 1).

### Characteristics of EQ-5D-3L and PHQ-9

With regard to the EQ-5D-3L results that respond to quality of life outcomes, it was shown that the main problems of university students are depression and moderate or severe anxiety (47.2%) and moderate or severe pain and discomfort (35.6%); while, personal self-care was the least frequent (4.6%). Regarding the visual analogue scale (EQ-VAS) of QoL, the average score obtained was 76.0 + 25.6 points (Table 2).

**Table 2.**
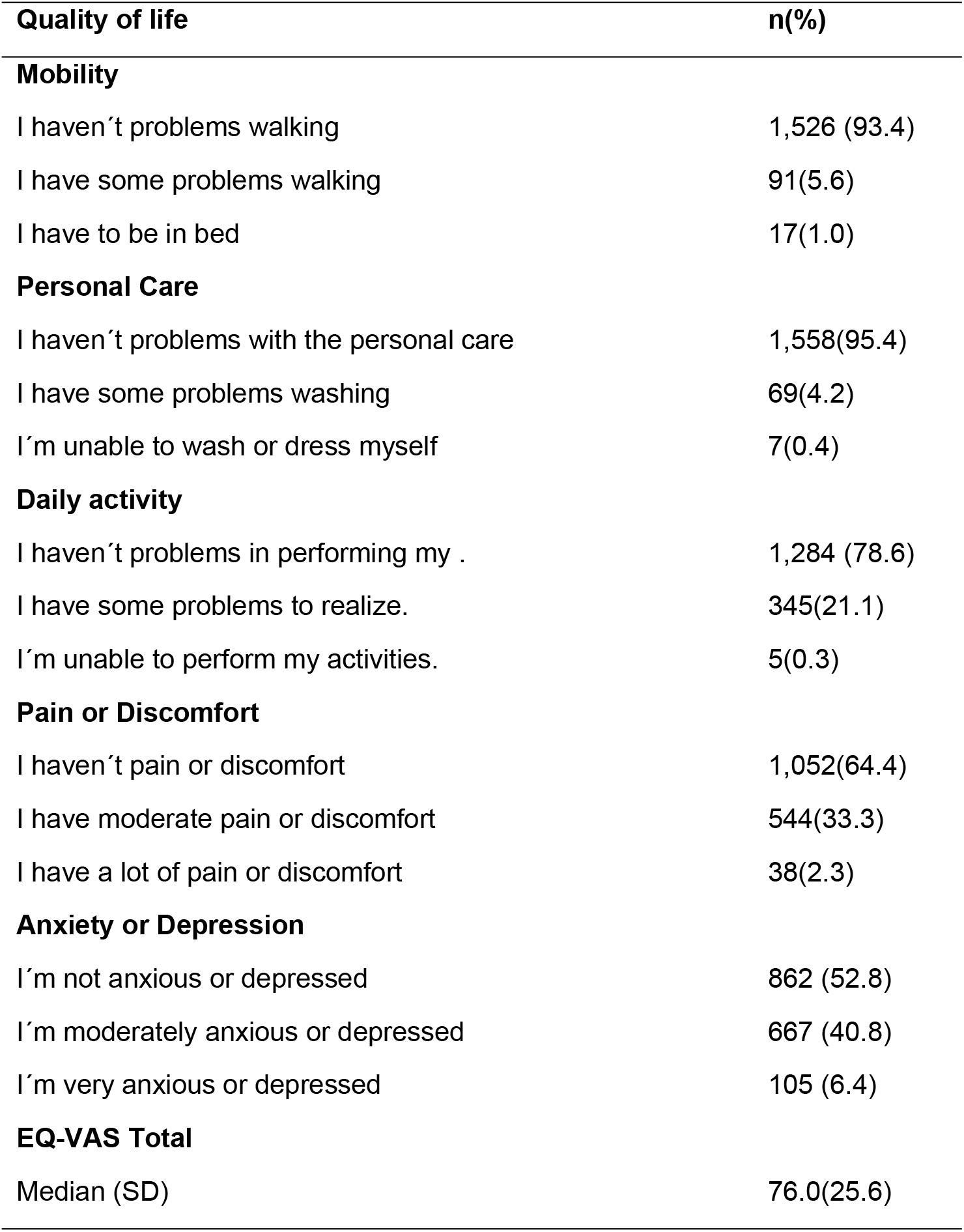
Quality of life among university students in Peru during the Covic-19 pandemic

| Quality of life | n(%) |
| --- | --- |
| **Mobility** |  |
| I haven't problems walking | 1,526 (93.4) |
| I have some problems walking | 91(5.6) |
| I have to be in bed | 17(1.0) |
| **Personal Care** |  |
| I haven't problems with the personal care | 1,558(95.4) |
| I have some problems washing | 69(4.2) |
| I'm unable to wash or dress myself | 7(0.4) |
| **Daily activity** |  |
| I haven't problems in performing my . | 1,284 (78.6) |
| I have some problems to realize. | 345(21.1) |
| I'm unable to perform my activities. | 5(0.3) |
| **Pain or Discomfort** |  |
| I haven't pain or discomfort | 1,052(64.4) |
| I have moderate pain or discomfort | 544(33.3) |
| I have a lot of pain or discomfort | 38(2.3) |
| **Anxiety or Depression** |  |
| I'm not anxious or depressed | 862 (52.8) |
| I'm moderately anxious or depressed | 667 (40.8) |
| I'm very anxious or depressed | 105 (6.4) |
| **EQ-VAS Total** |  |
| Median (SD) | 76.0(25.6) |

Likewise, through the PHQ-9 we found that the main depressive symptoms presented within a range of mild intensity (several days) to severe presence (almost every day) were feeling tired or low energy (69.9%), sleeping problems (62.1%), feeling down, depressed or hopeless (60.8%) and little interest or pleasure in doing things (59.06%); while, very rarely having ideas of death or suicide (16.3%). The level of depressive symptoms presented by university students was moderate to severe (28.9%) (Table 3).

**Table 3.** Depressive symptoms in university students in Peru during the Covid-19 pandemic

| **Depressive symptoms** | **n(%)** |
| --- | --- |
| **Low interest or pleasure in doing things** |  |
| Nothing | 669(40.94) |
| Several days | 572(35.0) |
| More than half the days | 191(11.7) |
| Almost every day | 202(12.36) |
| **You have felt down, depressed, or hopeless** |  |
| Nothing | 640(39.2) |
| Several days | 636(39.0) |
| More than half the days | 179(10.9) |
| Almost every day | 179(10.9) |
| **Difficulty sleeping or staying asleep or you have slept too much** |  |
| Nothing | 619(37.9) |
| Several days | 614(37.6) |
| More than half the days | 172(10.5) |
| Almost every day | 229(14.0) |
| **You have felt tired or low in energy** |  |
| Nothing | 492(30.1) |
| Several days | 727(44.5) |
| More than half the days | 212(13.0) |
| Almost every day | 203(12.4) |
| **Poor appetite or overeating** |  |
| Nothing | 746(45.7) |
| Several days | 545(33.4) |
| More than half the days | 183(11.2) |
| Almost every day | 160(9.7) |
| **You have felt bad about yourself or that you are a failure or that you have made a bad impression on your family** |  |
| Nothing | 893(54.7) |
| Several days | 470(28.8) |
| More than half the days | 154(9.4) |
| Almost every day | 117(7.1) |
| **You have had difficulty concentrating on things such as reading the newspaper or watching television** |  |
| Nothing | 799(48.9) |
| Several days | 518(31.7) |
| More than half the days | 177(10.8) |
| Almost every day | 140(8.6) |
| **Have you been moving or talking so slowly that other people might notice?** |  |
| Nothing | 1,026(62.8) |
| Several days | 408(25.0) |
| More than half the days | 120(7.3) |
| Almost every day | 80(4.9) |
| **You thought you'd be better off dead or thought you'd get hurt in some way** |  |
| Nothing | 1,367(83.7) |
| Several days | 152(9.3) |
| More than half the days | 65(4.0) |
| Almost every day | 50(3.0) |

**Table 3.** (Continued)
| **Depressive symptoms** | **n(%)** |
| --- | --- |
| **Depression** |  |
| Minimum | 686(42.0) |
| Slight | 475(29.1) |
| Moderated | 266(16.3) |
| Moderate-severe | 132(8.1) |
| Severe | 75(4.59) |

### Factors associated with EQ-VAS and PHQ-9

The univariate analysis reported that during the COVID-19 pandemic, sex and being a resident of the department of Lima or Piura is positively associated with the EQ-VAS (quality of life) score; while, age, occupation, residents of the department of Ayacucho, leaving home during quarantine, suffering decreased family income in quarantine, having a family member with chronic illness and depressive symptom scores are negatively associated. After adjusting the model, it was reported that age, occupation, flat of residence and leaving home in quarantine were no longer significant. In this model, QoL scores were significantly higher in men than in women (β=3.2, IC95%=1.1 to 5.4). Whereas, people who have suffered family economic decline during quarantine (β=-3.4, IC95%=-6.5 to −0.3), have family members with chronic diseases (β=-3.7, IC95%=-6.1 to −1.4) had significantly lower scores than people without these characteristics. Finally, it was reported that increasing 1 depressive symptom score reduced the quality of life score by 2 (β=-2.0, IC95%=-2.2 to −1.8) (Table 4).

**Table 4.** Factors associated with visual analogue scale of quality of life and PHQ-9 depression in university students in Peru during the COVID-19 pandemic

| Variables | Quality of life |  |  |  | Depressive symptoms |  |  |  |
| --- | --- | --- | --- | --- | --- | --- | --- | --- |
| | Crude<br>$\beta$ (IC95%) | p | Full<br>$\beta$ (IC95%) | p | Crude<br>$\beta$ (IC95%) | p | Full<br>$\beta$ (IC95%) | p |
| **Age** |  |  |  |  |  |  |  |  |
| 16-21 |  | REF |  |  |  | REF |  |  |
| 22-27 | -3.4 (-6.5;-0.3) | 0.032 | -2.3 (-5.0;0.4) | 0.101 | 0.51 (-0.2;1.2) | 0.170 | -0.1 (-0.6;0.6) | 0.986 |
| 28-65 | -0.29 (-3.2;2.6) | 0.846 | -2.0 (-4.8;0.8) | 0.164 | -0.76 (-1.5;-0.1) | 0.034 | -0.6 (-1.3;0.2) | 0.120 |
| **Sex** |  |  |  |  |  |  |  |  |
| Female |  | REF |  |  |  | REF |  |  |
| Male | 7.0 (4.5;9.4) | 0.000 | 3.2 (1.1;5.4) | 0.004 | -1.7 (-2.3; -1.1) | 0.000 | -0.7 (-1.3;-0.2) | 0.011 |
| **Marital Status** |  |  |  |  |  |  |  |  |
| Singer<br>/Separated/Widowed/Divorced |  | REF |  |  |  | REF |  |  |
| Married/Cohabitant | 1.7 (-1.2;4.5) | 0.239 | - | - | -1.1 (-1.7;-0.4) | 0.002 | -0.5 (-1.2;0.1) | 0.112 |
| **Ocupación** |  |  |  |  |  |  |  |  |
| Study and Work |  | REF |  |  |  | REF |  |  |
| Only Study | -2.0 (-4.5;0.5) | 0.118 | -0.7 (-3.0; 1.7) | 0.565 | 0.8 (0.2; 1.4) | 0.006 | 0.4 (-0.1;0.9) | 0.145 |
| **Departments of Peru** |  |  |  |  |  |  |  |  |
| Ancash |  | REF |  |  |  | REF |  |  |
| Ayacucho | -3.5 (-6.8;-0.2) | 0.040 | -2.1 (-4.8;0.8) | 0.151 | 0.9 (0.1;1.7) | 0.039 | 0.8 (0.1;1.5) | 0.022 |
| Lima | 4.0 (0.1;7.9) | 0.043 | 0.6 (-2.8;4.1) | 0.719 | -1.6 (-2.5;-0.6) | 0.001 | -0.7 (-1.5;0.2) | 0.116 |
| Piura | 3.6 (0.0;7.3) | 0.049 | 0.1 (-3.0; 3.3) | 0.933 | -1.6 (-2.5;-0.8) | 0.000 | -1.1 (-1.8;-0.3) | 0.005 |
| **He was out of the house during the quarantine** |  |  |  |  |  |  |  |  |
| Not |  | REF |  |  |  | REF |  |  |
| Yes | -3.2 (-5.7;-0.7) | 0.012 | -0.9 (-3.1;1.2) | 0.396 | 1.1 (0.5;1.7) | 0.000 | 0.7 (0.2; 1.2) | 0.006 |
| **Decrease in family income in quarantine** |  |  |  |  |  |  |  |  |
| Not |  | REF |  |  |  | REF |  |  |
| Yes | -8.1 (-11.6;-4.6) | 0.000 | -3.4 (-6.5;-0.3) | 0.034 | 1.9 (0.8;2.9) | 0.000 | 0.7 (-0.2;1.6) | 0.141 |
| Not |  | REF |  |  |  | REF |  |  |
| Yes | -2.1(-7.1;3.0) | 0.420 | - | - | 0.8 (-0.4;2.0) | 0.179 | 0.5 (-0.6;1.5) | 0.383 |
| **Family member with chronic illness** |  |  |  |  |  |  |  |  |
| Not |  | REF |  |  |  | REF |  |  |
| Yes | -8.7(-11.2; -6.1) | 0.000 | 3.7(-6.1; 1.4) | 0.002 | 2.4 (1.8 – 3.0) | 0.000 | 1.5 (1.0;2.1) | 0.000 |
| **Symptoms Depressives/Quality of life** | -2.1 (-2.3; -1.9) | 0.000 | -2.0 (-2.2; -1.8) | 0.000 | -0.1 (-0.2;-0.1) | 0.000 | -0.1 (-0.2;-0.1) | 0.000 |

On the other hand, factors associated with PHQ-9 scores (depressive symptoms) reported that all variables are significantly associated. However, after adjusting the model with all the variables, it was obtained that they remained significant: being residents of the department of Ayacucho, leaving home in quarantine and having a family member with chronic diseases (positive relationship); while, sex, being residents of the department of Piura and the quality of life score (negative relationship). Our positive relationship results with respect to the depressive symptom score refer that university students residing in Ayacucho presented higher scores compared to those residing in Ancash (β=0.8, IC95%=0.1 to 1.5), those who were quarantined compared to those who were not (β=0.7, IC95%=0.2 to 1.2) and having a relative with a chronic illness scored higher on depressive symptoms compared to those with no relatives with any illness (β=1.5, IC95%=1.0 to 2.1); the latter being the biggest problem. While, with respect to the negative relationship, men had lower scores than women (β=-0.7, IC95%=-1.3 to −0.2), being a resident of the department of Piura had lower scores than those living in Ancash (β=-1.1, IC95%=-1.8 to −0.3). Finally, the QoL score when decreasing 1 point of quality of life increases 0.1 points in depressive symptoms (β=-0.1, IC95%=-0.2 to −0.1) (Table 4).

## Discussion

University students in Peru during the COVID-19 pandemic presented an average EQ-VAS score of 76 points, almost half presented anxiety/depression and a third presented pain/discomfort. About a third presented moderate to severe depressive symptoms and more than two thirds felt tired or had low energy, had trouble sleeping, felt down, depressed or hopeless and presented a loss of interest or pleasure in doing things. QoL scores were significantly higher in men, while people who have suffered family economic decline during quarantine and have family members with chronic diseases had lower scores. Similarly, increasing 1 depressive symptom score reduced the QoL score by 2. With regard to factors associated with depressive symptoms, we found that being residents of the department of Ayacucho, leaving home in quarantine and having a family member with chronic diseases showed a positive relationship; while being a man, being a resident of the department of Piura and the quality of life score had a negative relationship.

In our population the dimensions of QoL most affected were anxiety/depression and pain/discomfort. This coincides with a study in Vietnamese university students during the COVID-19 pandemic, which reported a greater affect on the anxiety/depression and pain/discomfort dimensions of QoL (29). Another study of young Chinese adults reported similar effects on QoL dimensions during the pandemic (30). These two dimensions are often the most affected, possibly mourning the deaths of students’ relatives (31) and fear caused by overexposure to the media (8), which may have increased their anxiety and/or depression. Long months of confinement also resulted in constant exposure to stress and often manifested itself in pain and discomfort (32). (32) On the other hand, non-face-to-face education may have led to discomfort due to prolonged incorrect posture by students (33).

(33) University students who suffered a family economic downturn and had family members with chronic illnesses reported lower QoL scores. This is consistent with findings from other studies in China and Vietnam (30,34). Loss of economic income implies difficulty in accessing treatment, medication and living expenses, affecting quality of life (29,30,35). Men reported better QoL scores than women, similar to another recent finding (36), which could be due to the fact that men have a higher frequency of physical activity (37, 38) and less fear and distress generated by the VOC-19 pandemic (39, 40). We also found that, in view of the increase in the depressive symptom score, quality of life was reduced. Another study in Malaysia has found that depressive symptoms influence and impact the quality of life of university students (41).

Moderate and severe depressive symptoms occurred in almost one third of our population. As in a study with Chinese university students which reported a similar prevalence when assessing depressive symptoms during the Covid-19 pandemic (42). Other populations in China (43), Italy (44), Australia (45) and Mexico (46) report similar findings. The finding of the presence of depressive symptomatology in university students in Peru reinforces the idea of the effect on mental health during the Covid-19 pandemic.

This could be explained by several factors, for example, confinement as a measure to prevent the spread of Covid-19 and the impossibility of social interactions with peers is known to have a negative impact on mental health (4,47), sedentary behaviour adopted by students due to non-face-to-face education and financial insecurity for educational expenses have been associated with increased depressive symptoms in this population (48,49).

In our study two thirds of the university students felt tired or low in energy, had trouble sleeping, were depressed or hopeless, and had a loss of interest or pleasure in doing things. This coincides with studies in Saudi Arabia and China which reported a higher prevalence of those symptoms (50,51). Some studies have shown that these symptoms have been reported as associated reactions due to confinement and changes in habits that have had an impact on the mental health of the population (52).

Students residing in the department of Piura and Ayacucho reported lower and higher scores for depressive symptoms than residents of Ancash, respectively. This could be explained by the fact that during the months of evaluation of our study, the rate of COVID-19 infections and deaths in Piura had fallen, while in Ayacucho, a rural area of Peru, it was at its highest peak (53). The score of depressive symptoms was higher in women than in men; this is in line with a recent review study reporting gender differences in depression during COVID-19 (54). This is probably due to the fact that women are currently having more responsibilities within the work environment and during confinement the family burden (e.g. family caregivers) has increased, generating greater stress and depression(55,56). Students who left home during quarantine or had a family member with a chronic illness scored higher on depressive symptoms than those who did not have these characteristics, this finding could be due to the concern about catching and/or infecting at-risk family members during exposure to VOC-19 (57).

The findings in this study suggest the need to implement prevention and intervention strategies to mitigate problems in the quality of life and depressive symptoms of university students in Peru. The Peruvian government should continue to promote support such as educational grants for university students, it should also seek to gradually increase the number of places on such grants. The ministry of health, university institutions and the professional association of psychology should coordinate actions to provide telephone and online services as forms of strategies to assist in mental health and quality of life in this population (58). It is recommended that this study serve as a background for more complex studies and can also focus on evaluating interventions in the most affected groups, such as those with reduced income, with chronically ill family members and from rural Peru.

### Strengths and limitations

The strength of our study lies in the fact that it is the first study to report updated evidence on the state of quality of life and mental health of university students in Peru after the Covid-19 pandemic confinement. However, the study has some limitations. Due to the COVID-19 confinement, the sampling used was non-probabilistic which reduces the representativeness of the population findings, however, we reached an important sample size in different departments of Peru, which produces consistent evidence from university students, in addition the results obtained are similar to other studies conducted (29,42). Likewise, the instrument used (EQ-5D-3L) to evaluate the quality of life was not validated in Peru, however, this instrument has been used in other studies and populations in Peru (21,22), in addition it was translated into Spanish by the EuroQol Group and has been adapted in other countries and languages (20).

## Conclusions

In conclusion, quality of life has been affected in university students in Peru and they presented depressive symptoms during the COVID-19 pandemic. Specifically, more than half and almost half reported problems of pain/discomfort and anxiety/depression respectively. Specifically, more than half and almost half reported problems with pain/discomfort and anxiety/depression respectively. Men were found to have better QoL and those who had a lower income reported lower scores. Students from the department of Ayacucho, who left home in quarantine or had a family member with chronic illness were shown to be more likely to score higher on depressive symptoms while men, residents of the department of Piura and the quality of life score had a lower score compared to their reference categories.

